# Discordance in acute gastrointestinal toxicity between synchrotron-based proton and linac-based electron ultra-high dose rate irradiation

**DOI:** 10.1101/2024.09.04.611307

**Authors:** Kevin Liu, Uwe Titt, Nolan Esplen, Luke Connell, Elise Konradsson, Ming Yang, Xiaochun Wang, Takeshi Takaoka, Ziyi Li, Albert C. Koong, Devarati Mitra, Radhe Mohan, Billy W. Loo, Steven H. Lin, Emil Schüler

## Abstract

**Purpose:** Proton FLASH has been investigated using cyclotron and synchrocyclotron beamlines but not synchrotron beamlines. We evaluated the impact of dose rate (ultra-high [UHDR] vs. conventional [CONV]) and beam configuration (shoot-through [ST] vs. spread-out-Bragg-peak [SOBP]) on acute radiation-induced gastrointestinal toxicity (RIGIT) in mice. We also compared RIGIT between synchrotron-based protons and linac-based electrons with matched mean dose rates.

**Methods and Materials:** We administered abdominal irradiation (12-14 Gy single fraction) to female C57BL/6J mice with an 87 MeV synchrotron-based proton beamline (2 cm diameter field size as a lateral beam). Dose rates were 0.2 Gy/s (S-T pCONV), 0.3 Gy/s (SOBP pCONV), 150 Gy/s (S-T pFLASH), and 230 Gy/s (SOBP pFLASH). RIGIT was assessed by the jejunal regenerating crypt assay and survival. We also compared responses to proton [pFLASH and pCONV] with responses to electron CONV (eCONV, 0.4 Gy/s) and electron FLASH (eFLASH, 188-205 Gy/s).

**Results:** The number of regenerating jejunal crypts at each matched dose was lowest for pFLASH (similar between S-T and SOBP), greater and similar between pCONV (S-T and SOBP) and eCONV, and greatest for eFLASH. Correspondingly, mice that received pFLASH SOBP had the lowest survival rates (50% at 50 days), followed by pFLASH S-T (80%), and pCONV SOBP (90%), but 100% of mice receiving pCONV S-T survived (log-rank *P* = 0.047 for the four groups).

**Conclusions:** Our findings are consistent with an increase in RIGIT after synchrotron-based pFLASH versus pCONV. This negative proton-specific FLASH effect versus linac-based electron irradiation underscores the importance of understanding the physical and biological factors that will allow safe and effective clinical translation.

## INTRODUCTION

Ultra-high-dose rate (UHDR, >40 Gy/s) irradiation used in FLASH radiotherapy (RT) is of significant interest in the field of radiation oncology because of its promise for preferentially sparing normal tissue while maintaining isoeffective tumor response compared with conventional (CONV) dose rate (<1 Gy/s) irradiation (1–3). This phenomenon is termed the “FLASH effect.” Proton FLASH (pFLASH) RT has gained substantial momentum towards clinical translation over other UHDR modalities such as x-rays and electrons owing to its potential for improved tissue penetration and target conformality. The first human phase I clinical trial investigating the safety and feasibility of pFLASH in treating symptomatic bone metastases in the extremities (FAST-01) has now been completed, and a follow-up trial is currently recruiting patients that targets bone metastases in the thorax region (4–7).

To date, several studies have shown that pFLASH promotes normal tissue sparing following the irradiation of the skin, gastrointestinal (GI) tract, and brain when compared with pCONV (8–13). Meanwhile, other studies have been unable to demonstrate a significant pFLASH sparing effect and have even included reports of increased toxicity(6,14–17). The difficulties researchers have faced in reproducing the FLASH effect across different studies and institutions may reflect inconsistencies and uncertainties in the dosimetric conditions in UHDR beams, for which biological response may be differentially correlated with different physical beam parameters such as mean dose rate, instantaneous dose rate, number of pulses, temporal structure of the pulses or spills used (repetition rate), beam energy, the linear energy transfer (LET), field size, and others (3).

The current consensus is that the critical requirements for inducing the FLASH effect is sufficiently high dose being delivered at mean dose rates exceeding 40 Gy/s and in less than 200 ms (1,2,18). However, studies of electron-beam FLASH (eFLASH), in which beam parameters such as mean dose rate, pulse repetition frequency, pulse width, instantaneous dose rate, and dose per pulse are easily manipulated, have demonstrated that mean dose rate may not be the only factor at play, with the dose per pulse and instantaneous dose rate also reported as being key contributors to inducing and magnifying the FLASH effect (19–21). Comprehensive studies investigating the influence of temporal beam parameters have yet to be conducted with proton beamlines, likely because of difficulties in modifying beam parameters beyond just mean dose rate on proton accelerators. Proton beams used in RT are commonly produced by using one of three accelerator technologies: isochronous cyclotron (quasi-continuous, ns pulse structure), synchrocyclotron (µs pulse structure, comparable to that in electron beams), and synchrotrons (ms “spill” pulse structure), which result in three distinct temporal beam structures (22). Most investigations related to pFLASH have used cyclotron accelerators with double-scattered beams, but the effects of pFLASH have also been investigated with a single-scatter synchrocyclotron and with a cyclotron delivering a pencil-beam-scanning proton beam (8–11,13,15,16). However, to our knowledge, no studies have been published investigating differences in biological effects between pCONV and pFLASH *in vivo* when synchrotron accelerators are used.

The primary aim of the present study was to use a synchrotron-based pFLASH system to determine differences in the induction of radiation-induced normal tissue toxicity between mice treated with pCONV or pFLASH. A double-scattered 87-MeV peak energy beam was used, and acute GI toxicity was evaluated in both the shoot-through (S-T) and spread-out Bragg peak (SOBP) beam configurations (23,24). The proton irradiation response was then compared against previous work involving electron irradiation, with matched doses and mean dose rates (25). Previous studies with eCONV and eFLASH beams (19,25) showed that 12-14 Gy was the optimum dose range for evaluating differences in regenerating crypt counts between FLASH and CONV electron-based RT. This finding served as the basis for the range of doses used in the present study with protons.

## METHODS AND MATERIALS

### Irradiation setup

All irradiations were done with a Hitachi synchrotron (HITACHI, Ltd., Tokyo, Japan) under room air conditions, with a beamline modified to deliver pCONV and pFLASH with an energy of 87 MeV at the nozzle entrance and a range of 4.5 cm in water. Details of the beamline characteristics and commissioning are reported elsewhere (23,24) describing in detail its field flatness and symmetry for a circular field of 2-cm diameter, its ability to be shaped to deliver either a S-T or SOBP beam, and its ability to be adapted to provide UHDR in its current configuration. The LET of the S-T beam ranges from 0.9 to 1.1 keV/μm and the LET of the SOBP ranges from 1.25 to 2.8 keV/μm (23). Dosimetry performed on the proton beamline adhered to the standard described in the International Atomic Energy Agency TRS-398 report for absolute dose measurements (26) in both pFLASH and pCONV irradiation conditions, with dosimetric validation performed on the day of the experiment.

A custom-produced mouse immobilization setup was designed and seated on top of an acrylic phantom situated adjacent to the range compensator. The mouse cradle was designed to ensure stable and reproducible positioning of the mouse when stretched for lateral irradiations (Fig. 1A). Positioning uncertainty was determined by using mice matched for age, weight, and sex to the treated cohort (8-week-old female mice weighing approximately 18-20 g) by placing the mice (n=5) in the mouse cradle and imaged with a small-animal CT (SARRP, Xstrahl Inc., Swanee, GA, USA) (25). A 3D printed platform was then used to index the mouse cradle to the radiation field, with integrated attachments for delivery of anesthetics (isoflurane induction [3.5%, 450 mL/min, ∼2 min] and maintenance [1.5%, 150 mL/min, ∼ 3 min] with room air as carrier gas) situated adjacent to the range compensator (Fig. 1B, 1C). Figure 1C shows the schematic setup of the beamline for the SOBP configuration with the reference monitor chamber and beam profile monitor components upstream of the mouse platform for online dosimetry. The S-T configuration involves the removal of the energy filter in Figure 1C. Mice were irradiated in independent trials performed on separate days, with an average time between trials of 30 days. Trials were conducted to investigate beam configurations (S-T and SOBP) and assay types (crypt regeneration and survival), with each trial performed at least twice for a total of eight independent trials. pCONV and pFLASH irradiations within each trial were always performed on the same date.

**Figure 1.**
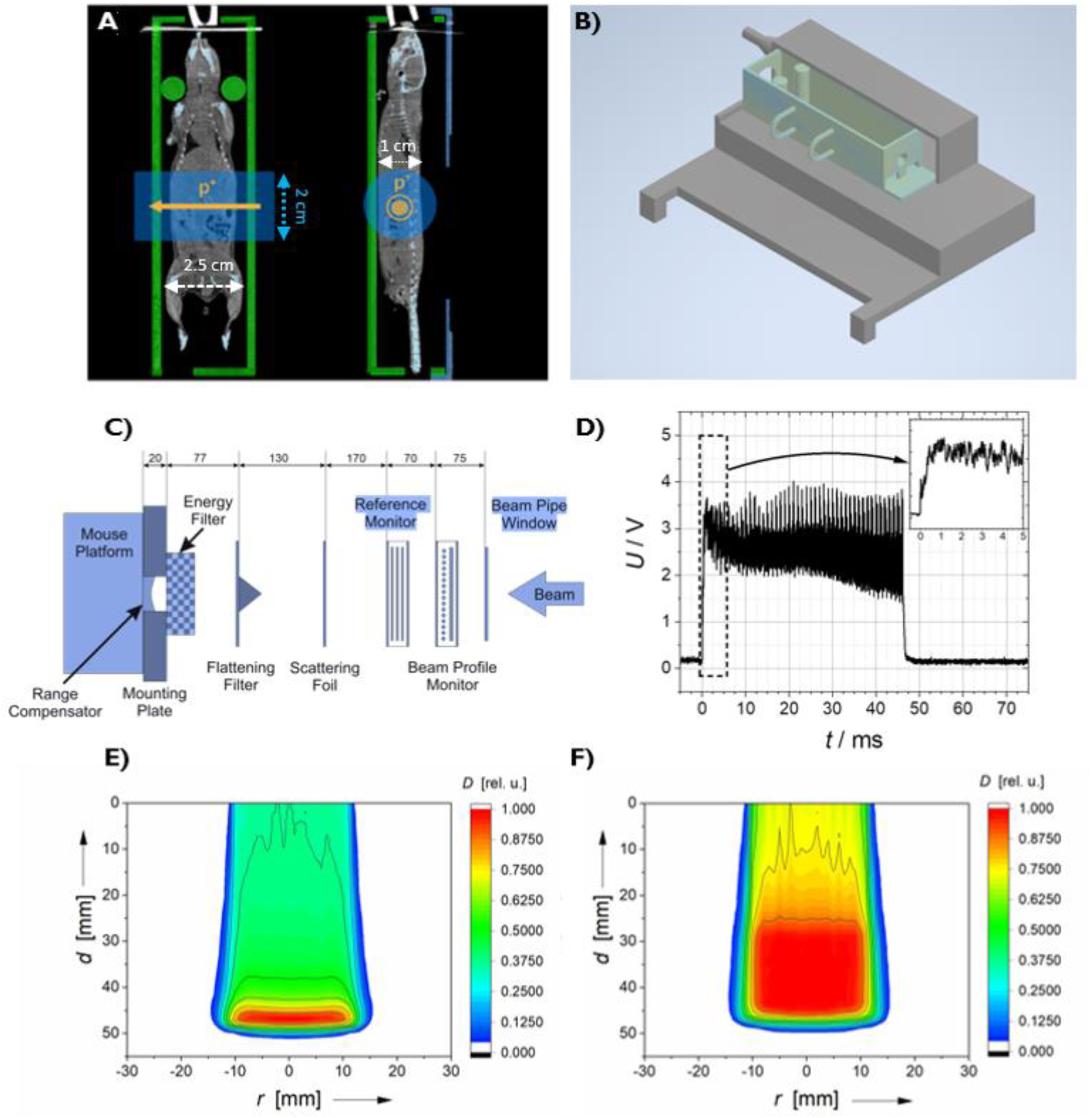
Detailed visualization of the experimental platform and beam configurations used for lateral proton (2-cm diameter) abdominal irradiation. (A) Coronal and sagittal CT slices of a mouse placed in the mouse cradle and then stretched for reproducible positioning, with columns to support the neck. The blue fields indicate where the mouse was irradiated (2-cm diameter), with the arrows indicating the direction of the proton beam. (B) The 3D printed design of the platform used to index the mouse cradle to the radiation field for the proton beamline. (C) The schematics of the beamline components used for spread-out-Bragg-peak (SOBP) proton irradiations, with distances labeled in units of mm. The energy filter is absent in the shoot-through (S-T) configuration. (D) Typical measured spill signal with high-frequency variations and low frequency changes measured from the reference chamber during a pFLASH dose delivery with the voltage (U) measured as a function of time, in ms, with the dose-rate being averaged from the measured signal over its spill duration. (E,F) The simulated depth distributions of the (E) S-T (without an energy filter) and (F) SOBP proton beams with a 20-mm modulation energy filter as a function of radial distance, *r*, from the central axis and of depth, *d*.

The irradiation setup was calibrated for pFLASH and pCONV conditions by using an Advanced Markus (PTW, Freiburg, Germany, SN 0294) ionization chamber connected to a DOSE1 electrometer (IBA, Hendon, Virginia, USA, SN 03-8673) for absolute dose measurements, with less than 1% recombination for mean dose rates up to 375 Gy/s as reported in a previous study of this synchrotron (24). Flatness/symmetry of the beam was evaluated by using dose-rate-independent Gafchromic film (Ashland Inc., Covington, Kentucky, USA) (27). The abdominal treatment dose was 12–14 Gy given in a single fraction, which corresponded to spill-times of 50-60 ms for the SOBP and 75-100 ms for the S-T pFLASH beams. Film was placed on the mouse cradle (entrance and exit) for each mouse irradiation to determine dosimetric consistency and to verify positioning. The time structure of each spill was measured with a Tektronix TDS3014B oscilloscope (Tektronix, Beaverton, OR, USA) during the beam-on period (Fig. 1D).The depth dose distributions for the S-T and SOBP proton beams are shown in Figures 1E and 1F.

pCONV irradiations were delivered at dose rates of 0.2 Gy/s for the S-T beam and 0.3 Gy/s for the SOBP beam, and pFLASH irradiations were delivered at dose rates of 150 Gy/s for the S-T beam and 230 Gy/s for the SOBP beam (Table 1). For pCONV irradiations, each spill had a duration of approximately 500 ms, with approximately a 2-second time interval between spills for passive scattering with an operation cycle of ∼2.5 seconds. For pFLASH irradiations, the radiofrequency extraction power pattern during the extraction time was adjusted to allow for higher charges to be deflected from their orbit into the extraction channel. This allows for all of the accelerated protons to be extracted within a single spill of 100 ms or less (23). No differences in the irradiation geometry was noted between pFLASH and pCONV irradiations for a given beam configuration (S-T or SOBP). pCONV irradiations involve multiple 500-ms spills with a smaller intra-spill dose rate and mean dose rate to deliver a given dose, whereas pFLASH irradiations involve a single spill (up to 100 ms) with a higher intra-spill and mean dose rates. Full control of beam deliveries was obtained by extracting the output signal from the reference monitor and feeding it to the amplifier of the dose monitor (Fig. 1C), threby allowing the user to stop the beam at a user-defined dose level, enabling partial spill deliveries in pFLASH and pCONV irradiation conditions(24).

**Table 1.** Experiment summary with measured dose, dose-rate, irradiation/spill time, and number of spills delivered for proton FLASH and CONV deliveries. The values represent the average and standard deviation or the range of values taken from n = 6-10 mice over the span of two independent trials.

| Treatment Condition | Group | Dose (Gy) | Mean Dose Rate (Gy/s) | Irradiation Time (s) | Number of Spills (Range) |
| --- | --- | --- | --- | --- | --- |
| Shoot-Through | FLASH | 12.02 ± 0.01 | 159 ± 13 | 0.076 ± 0.006 | 1 |
|  | FLASH | 13.00 ± 0.02 | 153 ± 13 | 0.085 ± 0.007 | 1 |
|  | FLASH | 13.97 ± 0.07 | 151 ± 14 | 0.094 ± 0.009 | 1 |
|  | CONV | 12.01 ± 0.02 | 0.23 ± 0.05 | 52.2 ± 10.4 | 30-32 |
|  | CONV | 13.01 ± 0.02 | 0.23 ± 0.05 | 57.6 ± 12.2 | 34-35 |
|  | CONV | 14.00 ± 0.01 | 0.23 ± 0.05 | 62.2 ± 12.9 | 37-38 |
| Spread-Out-Bragg-Peak | FLASH | 11.97 ± 0.02 | 233 ± 11 | 0.052 ± 0.002 | 1 |
|  | FLASH | 12.99 ± 0.01 | 229 ± 4.0 | 0.057 ± 0.001 | 1 |
|  | FLASH | 14.05 ± 0.02 | 230 ± 5.3 | 0.061 ± 0.001 | 1 |
|  | CONV | 12.01 ± 0.02 | 0.27 ± 0.01 | 44.0 ± 0.9 | 21-22 |
|  | CONV | 13.01 ± 0.01 | 0.27 ± 0.01 | 47.6 ± 1.6 | 23-24 |
|  | CONV | 14.02 ± 0.02 | 0.27 ± 0.01 | 51.6 ± 1.3 | 25-27 |

### Mouse handling

Eight-week-old female C57BL/6J mice from Jackson Laboratories (Bar Harbor, Maine, USA) were used. The mice were acclimatized for a minimum of 3 days before the experiment and had access to standard food and acidified water ad libitum. Five mice each were housed in ventilated cages in a 12/12-hour light/dark cycle. All mouse procedures in this work were approved by the Institutional Animal Care and Use Committee (IACUC) at our institution.

Radiation-induced GI toxicity (RIGIT) was evaluated by a jejunal crypt regeneration assay (n=6-10 mice/group). The protocol used followed that of previous investigations of the FLASH effect after total abdominal irradiation (19,25). At 89 ± 2 hours after irradiation, the mice were euthanized via CO_2_ inhalation followed by cervical dislocation to be within the timeframe for complete crypt loss followed by the formation of regenerating microcolonies (28–31). The jejunum from each mouse was collected, fixed in 10% neutral buffered formalin for 24 hours, washed with phosphate-buffered saline, and stored in 70% ethanol. Jejunum samples were then embedded in paraffin, and nine 3-μm thick transverse sections, separated by 5 mm along the length of the jejunum, were obtained and stained with hematoxylin and eosin. Only transverse sections with complete jejunal circumferences were included, and regenerating crypts were quantified by manual counting. A regenerating crypt was defined based on the following criteria: (1) basophilic structure along the circumferential edge, (2) U-shaped structure, and (3) multicellularity (at least 10 cells).

RIGIT was also assessed after pFLASH and pCONV by analyzing the survival of the mice over a period of 50 days. Mice were treated with a dose of 12 Gy under the following conditions (n=10/group): pFLASH S-T, pCONV S-T, pFLASH SOBP, pCONV SOBP, and SHAM. Throughout this period, mice were closely observed for indicators of acute toxicity such as weight loss, hunching, diarrhea, and changes in activity and behavior. Early euthanasia was initiated if the mice met the following IACUC criteria: (1) displaying moribund characteristics such as hunched posture, labored breathing, non-weight-bearing lameness or (2) experiencing weight less exceeding 30% of the baseline body weight.

### Statistical analysis

To assess normal tissue toxicity in terms of numbers of regenerating crypts in the pFLASH- and pCONV-treated groups, unpaired two-sample *t* tests were used to compare the number of regenerating crypts counted for different dose rate and beam configurations for a given dose. We also compared normal tissue effects between protons and electrons for the CONV and FLASH beamlines by using published data from a previous study (25). The electron FLASH (eFLASH) dose delivered was 12-14 Gy using an IntraOp Mobetron (IntraOp, Sunnyvale, CA, USA) with a mean dose rate of 188-205 Gy/s and 1.4–1.5 Gy per pulse (Table S1) measured using dosimeters and protocols described in past studies(32–37). The electron beam energy was 9 MeV and the field size used was 4 x 4 cm^2^(38,39). All statistical analyses were evaluated by using GraphPad Prism V.10 (La Jolla, CA, USA) and R (Auckland, NZ), with *P* values of <0.05 considered to indicate significant differences.

## RESULTS

### Dosimetry

The beam profiles measured on Gafchromic film placed on the mouse cradle for both pFLASH and pCONV in the S-T and SOBP beam configurations are shown in Figure 2. The dose profiles measured in the horizontal and vertical directions were found to be within 3% for both dose rate and beam configurations at the central 80% of the full-width-at-half-maximum (FWHM). The horizontal beam profile traces from the superior to inferior region of the mouse (Fig. 1A), and the vertical beam profile traces from the anterior to posterior region of the mouse. The penumbra in the anterior direction of the mouse was not measured on film in the beam as the 3D printed platform was designed to prevent steep dose gradients in the abdominal region during irradiations, as shown by the cut-off in the vertical profiles. The relative beam profiles and the dose measurements obtained with film (both entrance and exit) were found to be matched between mice treated at identical dose levels for pFLASH and pCONV within a trial.

**Figure 2.**
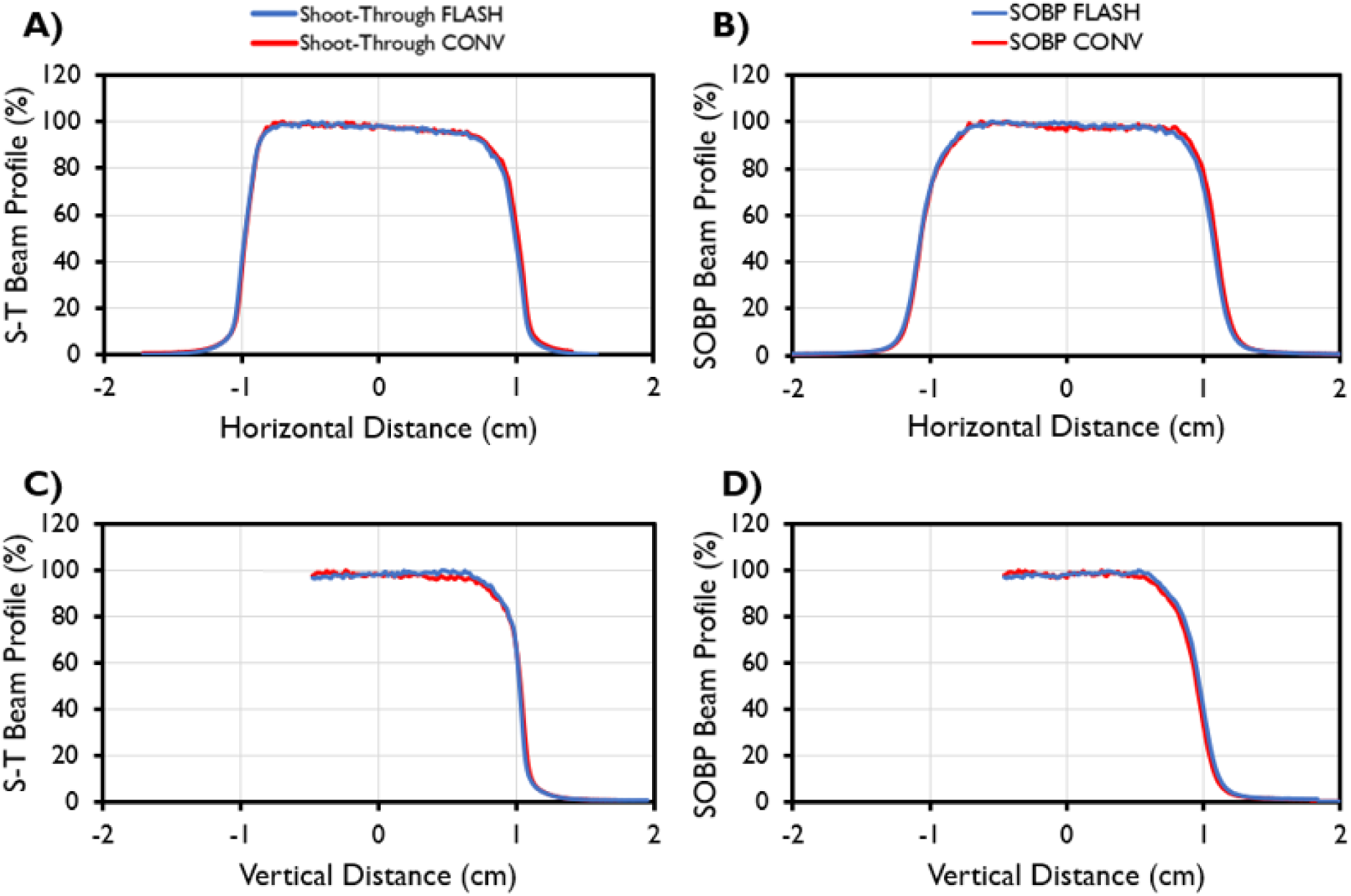
Dose distribution measured during *in vivo* experiments for both the shoot-through (S-T) and spread-out Bragg-peak (SOBP) proton beam configurations. (A-D) The beam profile measured in the horizontal direction for the (A) S-T and (B) SOBP and in the vertical direction for the (C) S-T and (D) SOBP measured at the mouse cradle by using Gafchromic EBT3 film for proton FLASH (blue) and proton CONV (red) RT. The horizontal beam profile begins from the mouse superior (left) to inferior (right). The vertical beam profile begins from the mouse anterior (left) to posterior (right).

### pFLASH produced greater acute radiation-induced GI toxicity than pCONV

RIGIT, evaluated with a regenerating crypt assay, was found to be significantly greater in mice treated with pFLASH than in those treated with pCONV, with the number of regenerating crypts being significantly lower at doses of 12 and 13 Gy for the S-T delivery and significantly lower at 12 Gy for the SOBP delivery (Fig. 3A, 3B). The numbers of regenerating crypts counted at 12 and 13 Gy for S-T and 12 Gy for SOBP were found to be 25%-60% lower for mice treated with pFLASH than for those treated with pCONV. Exposure to 14 Gy led to no significant differences in numbers of regenerating crypts between groups (*P*>0.05).

**Figure 3.**
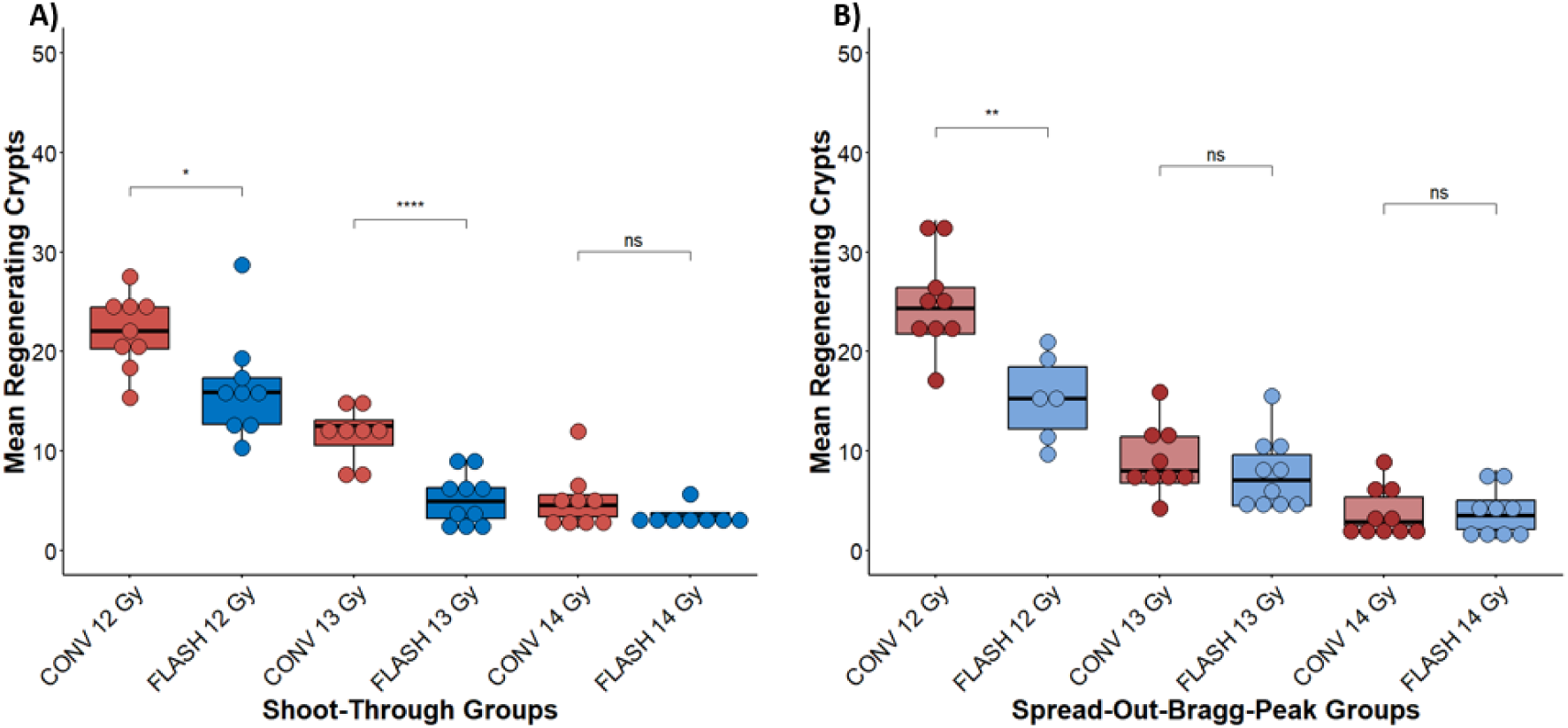
The mean numbers of regenerating crypts per circumference for mice receiving abdominal irradiation by either pCONV (0.2 / 0.3 Gy/s, Red) or pFLASH (150 / 230 Gy/s, Blue) to a dose of 12-14 Gy using (A) shoot-through (S-T) or (B) spread-out Bragg peak (SOBP) proton beam configurations. pFLASH-treated groups were compared with pCONV-treated groups with an unpaired parametric *t* test.\**P*< 0.05; \*\**P*< 0.01; \*\*\**P*< 0.001; \*\*\*\**P*< 0.0001. For both S-T and SOBP, the mean numbers of regenerating crypts were significantly fewer at 12 and 13 Gy, but no different at 14 Gy, for pFLASH vs pCONV.

### eFLASH produced less acute RIGIT than pFLASH, but no differences were found in eCONV vs pCONV

The mean numbers of regenerating jejunal crypts counted under identical dose and dose rate conditions were compared for mice treated under different beam modalities (protons [with SOBP and S-T] vs electrons; Fig. 4). No differences were found in the numbers of regenerating crypts in mice treated with pCONV for both the S-T and SOBP configurations vs eCONV at doses of 12–14 Gy. However, under FLASH conditions, mice treated with electrons (eFLASH) exhibited the opposite response from protons, with significantly lower GI toxicity (*P*<0.01) relative to both S-T and SOBP pFLASH conditions at doses of 12–14 Gy. The numbers of regenerating crypts in mice treated with eFLASH were 2-5 times higher than in mice treated with pFLASH. No differences were found in numbers of regenerating crypts between the S-T and SOBP configurations in either pFLASH or pCONV.

**Figure 4.**
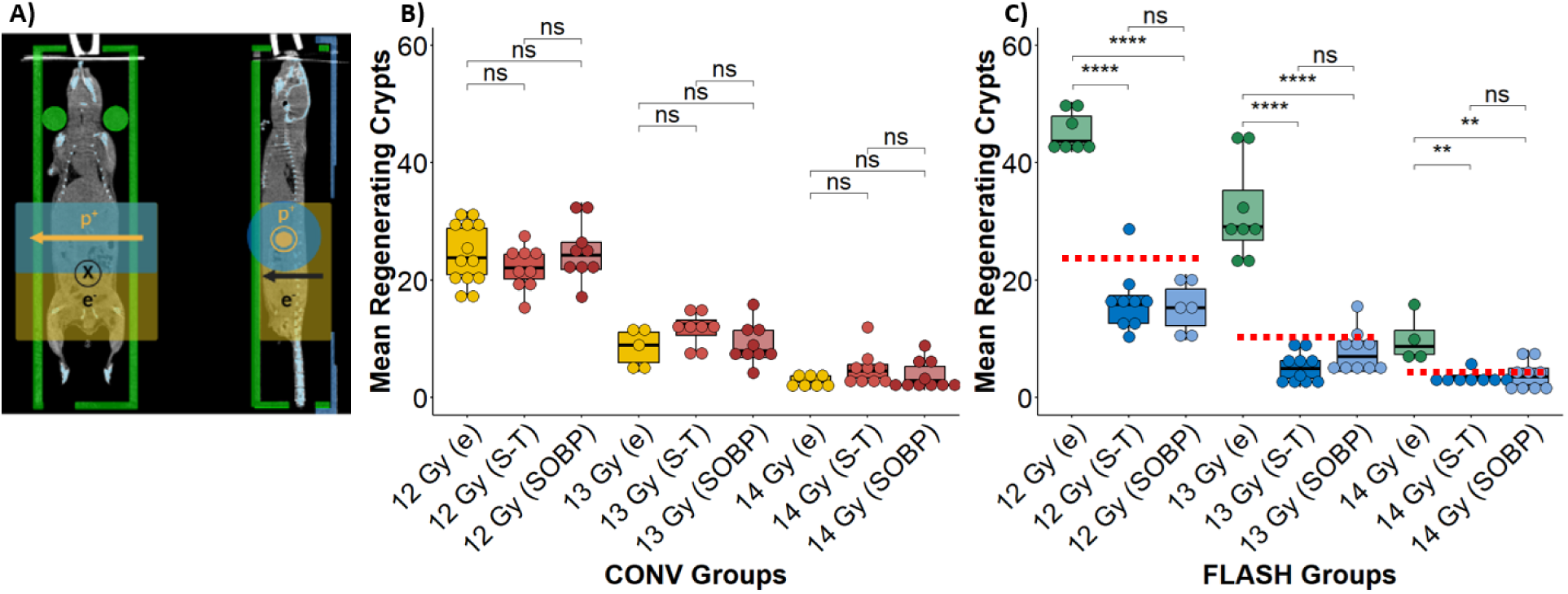
(A) Coronal and sagittal CT slices of a mouse placed in the mouse cradle, with columns supporting the neck, for lateral proton (2-cm diameter) and posterior-anterior electron (4 x 4 cm^2^) abdominal irradiation. (B,C) The mean numbers of regenerating crypts per circumference for mice receiving abdominal irradiation by shoot-through (S-T, 0.2 Gy/s pCONV, 150 Gy/s pFLASH) and spread-out Bragg peak (SOBP, 0.3 Gy/s pCONV, 230 Gy/s pFLASH) for (B) CONV or (C) FLASH to a dose of 12-14 Gy for an 87 MeV proton beam and a 9 MeV electron beam (0.4 Gy/s eCONV, 187-206 Gy/s eFLASH). The red-dotted line in (C) correspond to the mean number of regenerating crypts in the corresponding CONV groups, for comparison. Groups treated with CONV or FLASH were compared using an unpaired parametric *t* test. \**P*< 0.05; \*\**P*< 0.01; \*\*\**P*< 0.001; \*\*\*\**P*< 0.0001. Under CONV, no difference in the mean number of regenerating crypts were found between mice treated with protons or electrons. Under FLASH, the mean number of regenerating crypts was higher in mice treated with electrons than with protons irrespective of S-T or SOBP. Under both pFLASH and pCONV, no difference in the mean number of regenerating crypts were found between S-T and SOBP. [Electron data from Valdez Zayas et al., 2023.](40)

### pFLASH produced higher mortality than pCONV in both Shoot-Through and Spread-Out Bragg Peak beam configurations, with higher mortality in the Spread-Out Bragg-Peak condition

In evaluating survival over 50 days, mice that were treated with S-T under pCONV and pFLASH conditions had lower mortality rates than mice treated with SOBP under both conditions. Relative survival rates were 100% for the S-T / pCONV to 12 Gy vs 80% for the S-T / pFLASH group. Log-rank tests showed no significant differences in survival between pFLASH, pCONV, and SHAM conditions (*P*>0.05) for mice treated with S-T. Survival rates were 90% for mice treated with SOBP / pCONV to 12 Gy vs 50% for those treated with SOBP / pFLASH. Log-rank tests showed a survival difference between pFLASH, pCONV, and SHAM for mice treated with SOBP (*P*<0.05). Survival rates were also different for mice treated with S-T pFLASH, SOBP pFLASH, and SHAM using a log-rank test (*P*<0.05) (Fig. 5).

**Figure 5.**
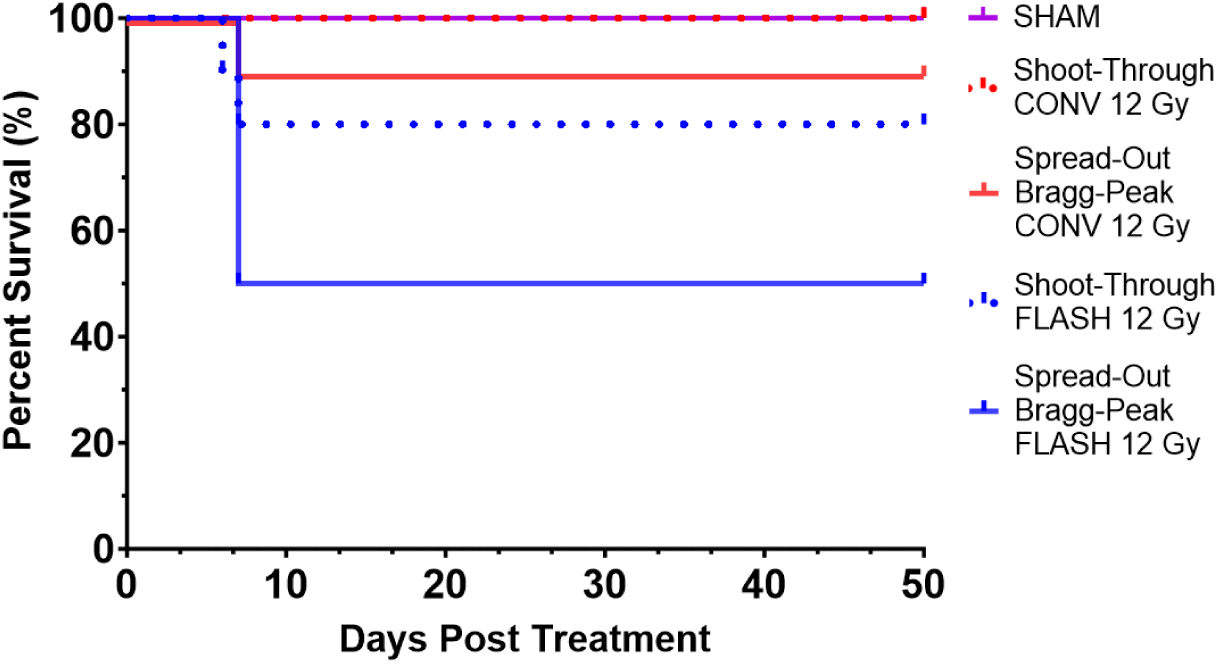
Kaplan-Meier plots of survival as a function of days after treatment (up to 50 days) with 12 Gy of pFLASH or pCONV to the abdomen with the shoot-through (S-T) or spread-out Bragg peak (SOBP) beam configurations, with a SHAM (unirradiated) group for comparison (n=10/group). pFLASH led to decreased survival relative to pCONV for both S-T and SOBP, with higher mortality after SOBP than after S-T. Mice treated with pFLASH SOBP had the lowest survival (50% survival), followed by pFLASH S-T and pCONV SOBP (80-90%), while 100% of mice receiving SHAM treatment and pCONV S-T survived.

## DISCUSSION

FLASH is rapidly developing towards clinical translation, with pFLASH currently being the only accessible modality capable of reaching deep-seated tumors while maintaining dose rates of >40 Gy/s, with potential for conformal dose distributions (41). Although preclinical research is crucial for determining if the FLASH effect can be translated to humans, results have been conflicting regarding the magnitude and reproducibility of the FLASH effect in preclinical settings (3). In this study, we conducted the first investigation of CONV and FLASH irradiation using synchrotron-accelerated protons on a clinical system and investigated the magnitude of RIGIT, with mice irradiated in the S-T or in the SOBP beam configurations. Every effort was taken to ensure robustness and reproducibility in our irradiations and setups. The system was calibrated on each day of irradiation by using previously established calibration protocols (23,24,26). Online dosimetry was performed with a repurposed reference monitor chamber and beam profile monitor for beam control and beam profile measurements (Fig. 1C) (23). Spatial dose distribution and dosimetric consistency were further validated by using film dosimetry at the beam entrance and exit of the mouse cradle for each mouse that was treated. Use of a 3D printed immobilization jig enabled reproducible mouse positioning within the weight range used to an accuracy of better than 1.5 mm as determined by CT. The jig was indexed to the radiation beam by using an indexing platform to ensure reproducible alignment to the beam. Furthermore, each treatment condition was independently reproduced on a different date to add further robustness to our approach and analysis.

A total of eight separate trials was conducted, with two independent trials for each treatment condition (S-T or SOBP) and for each assay (crypt regeneration or survival), with FLASH and CONV always delivered on the same date for the same beam configuration. The dose variation within and between trials was <1%, and the dose variation between CONV and FLASH deliveries was within 1% based on the dosimetric protocols. Nine 5-mm sections of the small intestines were extracted and analyzed to enhance sampling of the response along the entirety of the jejunum. No significant difference between sections was found, confirming homogeneous dose delivery along the analyzed jejunum. Also, good consistency in the biological data acquired was confirmed between the trials (allowing the data to be combined in the final analysis).

Independent of treatment condition, pFLASH was found to consistently induce greater toxicity than pCONV. This finding stands in stark contrast to previously published pFLASH studies using cyclotron or synchrocyclotron accelerators (9,11,42,43) regarding RIGIT, and to other studies of eFLASH (19,20,25,44,45), in which FLASH was shown to induce significantly less RIGIT than CONV with the assays used here. To our knowledge, only one other study has investigated eFLASH and pFLASH together (46). In that study, mice treated with UHDR whole-brain irradiation exhibited neurocognitive function similar to that in SHAM-treated mice regardless of radiation type, but mice treated with whole-brain irradiation at CONV dose rates showed significant cognitive decrements. Interestingly, greater differences in cognitive sparing between FLASH- and CONV-treated mice were found in the electron cohort than in the proton cohort. That study also confirmed complete tumor response in mice treated with both FLASH and CONV for both protons and electrons (46).

In a direct comparison between mice treated in the current study with protons and those treated with electrons in a prior study conducted by our group, which employed similar dose and dose rate conditions, the number of regenerating crypts did not differ between eCONV and pCONV(25). This equivalent level of jejunal injury was seen despite different field sizes (4×4 cm^2^ field size for eCONV and 2-cm diameter field size for pCONV), likely reflecting similar exposure of jejunal tissue to similar radiation doses and dose rates. However, eFLASH showed significant sparing of jejunal tissue when compared with eCONV; this is in contrast to pFLASH in the present work that showed a significant increase in jejunal injury relative to pCONV.

In the current study, the mean dose rates of the S-T and SOBP beamlines were not matched under UHDR conditions: the mean dose rate was higher for the SOBP proton beam. Despite this difference in mean dose rate, the response between S-T and SOBP were the same. A prior study investigating pFLASH (110 Gy/s) vs pCONV (0.8 Gy/s) under S-T and SOBP beam configurations, at matched mean dose rates between the beam configurations, also found no significant difference in the number of regenerating crypts between the S-T and SOBP beam configuration (9). The lack of difference in regenerating crypts in mice treated using S-T and SOBP, regardless of the difference in mean dose rate conditions across these studies, may reflect their comparable RBE at the total dose delivered and the assays used. A previous study investigating the RBE of 85 MeV proton beams through the use of a jejunal crypt cell assay reported an RBE of 1.08 at a delivered dose of 13.5 Gy (47), with other published values of RBE being of similar magnitude for the 12-14 Gy dose range (48–50). When we used survival as an assay to evaluate long-term GI toxicity for mice treated with S-T and SOBP for both pFLASH and pCONV at 12 Gy, we found that despite the same physical dose being delivered to the abdomen for mice treated, mice that were irradiated with the SOBP beam had shorter survival than those treated with the S-T beam, regardless of the mean dose rate. In a past study (25), actuarial survival of mice treated with electrons showed 100% and 90% survival rates in both FLASH and CONV at total abdominal doses exceeding 12 Gy. It is currently unknown why mice treated with a larger field size and a higher dose with electrons demonstrate a greater survival rate than mice treated with protons despite comparable dose rate conditions. However, this discrepancy may reflect confounding experimental variables which included differences in beam energy (which influences the depth dose and lateral dose distribution), differences in the output and temporal structures between the two different beam types, variability in mouse handling and transportation, batch differences from the vendor, and gut microbiome of the mice between studies (51).

This study is one of the few published to date indicating that mice treated with pFLASH experienced greater normal tissue toxicity than those treated with pCONV. Although the reasons behind this differential effect are unknown, one potential explanation may be the substantially lower instantaneous dose rate delivered from this synchrotron accelerator compared with cyclotron-based accelerators and eFLASH linear accelerators (8). For pFLASH, normal tissue sparing has been found to be sensitive to dose rate, time structure, and total irradiation time (52,53). In the current study, we used comparable mean dose rates but had instantaneous dose rates that were approximately 10 times lower than what is delivered from cyclotron-based proton accelerators, and several orders of magnitude lower than what is delivered from electron linear accelerators used for eFLASH (14,19,25,54). In our synchrotron system, the instantaneous dose rate of a UHDR proton delivery varies depending on the spill time: the ranges of spill time were 50-60 ms for SOBP and 70-100 ms for S-T, with lower instantaneous dose rates measured at longer spill times (24). Desoute this low instantaneous dose rate, the current work is not the only study to report greater normal tissue toxicity using pFLASH (a negative FLASH effect) when using proton-based accelerators.

A study conducted at Massachusetts General Hospital (MGH) used a cyclotron accelerator with 120 Gy/s for the pFLASH beam. The initial pilot study (43) indicated a potential tissue sparing effect in mice treated with pFLASH relative to those treated with pCONV; however, a subsequent study with larger numbers of mice and different irradiation schemes demonstrated greater normal tissue toxicity in the pFLASH-treated group (15). Both of the MGH studies used a beam structure similar to that used by Diffenderfer et al (42), who reported a positive FLASH effect with the same type of cyclotron system (IBA, Louvain-La-Neuve, Belgium), same radiofrequency (106 MHz), and similar pulse length (2–3 ns) (15), with the difference being the field size (16 x 12 mm^2^ elliptical field used by Zhang et al (15,43) vs 20 x 20 mm^2^ square field used by Diffenderfer et al (42)). Likewise, a separate study from Montefiore Einstein Cancer Center used a 250 MeV Varian ProBeam system (Varian Medical Systems, a Siemens Healthineers company, Palo Alto, California, USA) that involves an isochronous cyclotron that delivers proton RT in pencil-beam scanning mode, with field-averaged dose rates of 0.6 Gy/s for pCONV and 80–100 Gy/s for pFLASH (16). That group also reported a significant decrease (by 40%) in the survival of mice treated with pFLASH at 14 Gy. They also found no significant differences in the measured histologic endpoints such as intestinal crypts, EdU+ cells/crypt, and Olfactomedin4+ intestinal stem cells at delivered doses ranging from 12 to 15 Gy for pFLASH and pCONV. In these two studies and ours, pFLASH beams produced by different accelerator types and modalities (synchrotron vs cyclotron or double-scatter vs pencil beam scanning) demonstrated significantly greater normal tissue toxicity after abdominal FLASH irradiation. Currently, it is unknown as to why using protons at UHDR would be more toxic to normal tissue than protons delivered at CONV dose rates. Also, it remains unknown as to why data are conflicting between institutions despite the use of similar beam structures and beam configurations. Future work is warranted to investigate potential reasons behind the differences in biological response on pFLASH beamlines.

## CONCLUSIONS

The differential effects of synchrotron-produced proton irradiation was investigated under FLASH and CONV conditions on RIGIT after abdominal irradiation of female C57BL/6J mice. Mice irradiated under pFLASH conditions exhibited significantly fewer regenerating crypts than those irradiated under pCONV conditions for both S-T and SOBP proton beams. No statistically-significant differences were found in the regenerating crypt assay of mice treated to the same dose in the S-T vs the SOBP region of the proton beam irrespective of pFLASH or pCONV conditions. Correspondingly, survival analysis revealed greater mortality in mice treated using pFLASH than in those treated using pCONV, and higher mortality rates for the SOBP vs the S-T beam. In comparing the response to protons and electrons under similar CONV and FLASH conditions, we found no significant differences between mice treated with protons and electrons under CONV conditions—however, eFLASH resulted in significant crypt sparing and increase in survival compared with eCONV-irradiated mice — a response that was opposite to that observed using protons. These findings indicate that induction of the FLASH effect may depend not only on mean dose rate but also on the modality, radiation type, and other beam parameters. Future studies will investigate if increased acute toxicity is also evident in other tissues or whether FLASH and CONV irradiation produce different long-term effects on the GI tract.

## Acknowledgements

We thank Christine F. Wogan, MS, ELS, of MD Anderson’s Division of Radiation Oncology, for editorial contributions to this article.

## Funding

Research reported in this publication was supported in part by the National Cancer Institute of the National Institutes of Health under R01CA266673 and P30 CA016672; by the University Cancer Foundation via the Institutional Research Grant program at MD Anderson Cancer Center; by The University of Texas MD Anderson Cancer Center, Division of Radiation Oncology; by The University of Texas MD Anderson Cancer Center UTHealth Graduate School of Biomedical Sciences Dr. John J. Kopchick Fellowship; and by UTHealth Innovation for Cancer Prevention Research Training Program Pre-doctoral Fellowship (Cancer Prevention and Research Institute of Texas grant RP210042). The content is solely the responsibility of the authors and does not necessarily represent the official views of the National Institutes of Health or the Cancer Prevention and Research Institute of Texas.

## Conflicts of Interest

BWL is a cofounder and board member of TibaRay. SHL reports receiving grants from Nektar Therapeutics, Beyond Spring Pharmaceuticals, and STCube Pharmaceuticals; serving on the advisory board of AstraZeneca and Creatv Microtech; consulting fees from XRAD Therapeutics; and stock options in SEEK Diagnostics (co-founder). TT is currently employed by Hitachi Ltd.

## Data Availability

Research data are stored in an institutional repository and will be shared upon request to the corresponding author.

## Supplementary Data

**Table S1.** Beam parameters used for treating mice with electron irradiation on the IntraOp Mobetron.

| Group | Dose, Gy | Dose per Pulse, Gy | Mean Dose Rate, Gy/s | Irradiation Time, s | Intrapulse Dose Rate, MGy/s | Pulse Width, μs | Pulse Repetition Frequency, Hz | Number of Pulses/MU |
| --- | --- | --- | --- | --- | --- | --- | --- | --- |
| FLASH 12 Gy | 11.9 ± 0.3 | 1.49 ± 0.03 | 204.7 ± 4.2 | 0.058 ± 0.000 | 1.24 ± 0.03 | 1.2 | 120 | 8 |
| FLASH 13 Gy | 12.8 ± 0.1 | 1.42 ± 0.02 | 192.3 ± 2.1 | 0.067 ± 0.000 | 1.19 ± 0.01 | 1.2 | 120 | 9 |
| FLASH 14 Gy | 14.1 ± 0.1 | 1.41 ± 0.01 | 188.2 ± 1.9 | 0.075 ± 0.000 | 1.18 ± 0.01 | 1.2 | 120 | 10 |
| CONV 12 Gy | 12.0 ± 0.3 | 0.014 ± 0.000 | 0.4 ± 0.0 | 27.2 ± 0.6 | 0.01 ± 0.00 | 1.2 | 30 | 900 |
| CONV 13 Gy | 13.0 ± 0.2 | 0.014 ± 0.000 | 0.4 ± 0.0 | 29.4 ± 0.4 | 0.01 ± 0.00 | 1.2 | 30 | 975 |
| CONV 14 Gy | 14.2 ± 0.2 | 0.014 ± 0.000 | 0.4 ± 0.0 | 32.2 ± 0.4 | 0.01 ± 0.00 | 1.2 | 30 | 1050 |

## REFERENCES

1. Favaudon V., L. Caplier, V. Monceau, et al. Ultrahigh dose-rate flash irradiation increases the differential response between normal and tumor tissue in mice. Science Translational Medicine 2014;6.

2. Friedl A. A., K. M. Prise, K. T. Butterworth, et al. Radiobiology of the flash effect. Medical Physics 2022;49:1993–2013.

3. Schuler E., M. Acharya, P. Montay-Gruel, et al. Ultra-high dose rate electron beams and the flash effect: From preclinical evidence to a new radiotherapy paradigm. Med Phys 2022;49:2082–2095.

4. Daugherty EC, AE Mascia, MGB Sertorio, et al. Fast-01: Results of the first-in-human study of proton flash radiotherapy. International Journal of Radiation Oncology, Biology, Physics 2022;114:S4.

5. Daugherty E. C., A. Mascia, Y. Zhang, et al. Flash radiotherapy for the treatment of symptomatic bone metastases (fast-01): Protocol for the first prospective feasibility study. JMIR Res Protoc 2023;12:e41812.

6. Mascia A. E., E. C. Daugherty, Y. Zhang, et al. Proton flash radiotherapy for the treatment of symptomatic bone metastases: The fast-01 nonrandomized trial. JAMA Oncol 2023;9:62–69.

7. Daugherty E. C., Y. Zhang, Z. Xiao, et al. Flash radiotherapy for the treatment of symptomatic bone metastases in the thorax (fast-02): Protocol for a prospective study of a novel radiotherapy approach. Radiat Oncol 2024;19:34.

8. Diffenderfer E. S., B. S. Sorensen, A. Mazal, et al. The current status of preclinical proton flash radiation and future directions. Med Phys 2022;49:2039–2054.

9. Kim M. M., Verginadis, II, D. Goia, et al. Comparison of flash proton entrance and the spread-out bragg peak dose regions in the sparing of mouse intestinal crypts and in a pancreatic tumor model. Cancers (Basel) 2021;13.

10. Cunningham S., S. McCauley, K. Vairamani, et al. Flash proton pencil beam scanning irradiation minimizes radiation-induced leg contracture and skin toxicity in mice. Cancers (Basel) 2021;13.

11. Evans T., J. Cooley, M. Wagner, et al. Demonstration of the flash effect within the spread-out bragg peak after abdominal irradiation of mice. Int J Part Ther 2022;8:68–75.

12. Iturri L., A. Bertho, C. Lamirault, et al. Proton flash radiation therapy and immune infiltration: Evaluation in an orthotopic glioma rat model. Int J Radiat Oncol Biol Phys 2023;116:655–665.

13. Velalopoulou A., I. V. Karagounis, G. M. Cramer, et al. Flash proton radiotherapy spares normal epithelial and mesenchymal tissues while preserving sarcoma response. Cancer Res 2021;81:4808–4821.

14. Beyreuther E., M. Brand, S. Hans, et al. Feasibility of proton flash effect tested by zebrafish embryo irradiation. Radiotherapy and Oncology 2019;139:46–50.

15. Zhang Q., L. E. Gerweck, E. Cascio, et al. Absence of tissue-sparing effects in partial proton flash irradiation in murine intestine. Cancers (Basel) 2023;15.

16. Bell B.I., Velten, C., Pennock, M., Kang, M., Tanaka, K.E., Selvaraj, B., Bookbinder, A., Koba, W., Vercellino, J., English, J., Małachowska, B., Pandey, S., Duddempudi, P.K., Yang, Y., Shajahan, S., Hasan, S., Choi, J.I., Simone, C.B., Yang, W., Tomé, W.A., Lin, H., Guha, C. Pencil beam scanned proton flash radiotherapy increases acute lethality. (Submitted) 2024.

17. Kacem H., S. Psoroulas, G. Boivin, et al. Comparing radiolytic production of h(2)o(2) and development of zebrafish embryos after ultra high dose rate exposure with electron and transmission proton beams. Radiother Oncol 2022;175:197–202.

18. Kacem H., A. Almeida, N. Cherbuin, et al. Understanding the flash effect to unravel the potential of ultra-high dose rate irradiation. Int J Radiat Biol 2022;98:506–516.

19. Kevin Liu Trey Waldrop, Edgardo Aguilar, Nefetiti Mims, Denae Neill, Abagail Delahoussaye, Ziyi Li, David Swanson, Steven H Lin, Albert C Koong, Cullen M Taniguchi, Billy W Loo Jr, Devarati Mitra, Emil Schueler. Redefining flash rt: The impact of mean dose rate and dose per pulse in the gastrointestinal tract. bioRxiv 2024.

20. Ruan Jia-Ling, Carl Lee, Shari Wouters, et al. Irradiation at ultra-high (flash) dose rates reduces acute normal tissue toxicity in the mouse gastrointestinal system. International Journal of Radiation Oncology* Biology* Physics 2021;111:1250–1261.

21. Montay-Gruel P., M. M. Acharya, P. Goncalves Jorge, et al. Hypofractionated flash-rt as an effective treatment against glioblastoma that reduces neurocognitive side effects in mice. Clin Cancer Res 2021;27:775–784.

22. Jolly S., H. Owen, M. Schippers, et al. Technical challenges for flash proton therapy. Phys Med 2020;78:71–82.

23. Titt U., M. Yang, X. Wang, et al. Design and validation of a synchrotron proton beam line for flash radiotherapy preclinical research experiments. Medical physics 2022;49:497–509.

24. Yang M., X. Wang, F. Guan, et al. Adaptation and dosimetric commissioning of a synchrotron-based proton beamline for flash experiments. Physics in medicine and biology 2022;67.

25. Valdés Zayas A., N. Kumari, K. Liu, et al. Independent reproduction of the flash effect on the gastrointestinal tract: A multi-institutional comparative study. Cancers 2023;15.

26. Andreo P Burns DT, Hohlfeld K, Huq MS, Kanai T, Laitano F, Smyth V, Vynckier S. Iaea trs-398–absorbed dose determination in external beam radiotherapy: An international code of practice for dosimetry based on standards of absorbed dose to water. International atomic energy agency. International Atomic Energy Agency 2000;18:35–36.

27. Liu K Jorge PG, Tailor R, Moeckli R, Schüler E. Comprehensive evaluation and new recommendations in the use of gafchromic ebt3 film Med Phys 2023.

28. Hagemann R. F., C. P. Sigdestad, S. Lesher. Intestinal crypt survival and total and per crypt levels of proliferative cellularity following irradiation: Fractionated x-ray exposures. Radiat Res 1971;47:149–58.

29. Potten C. S. The significance of spontaneous and induced apoptosis in the gastrointestinal tract of mice. Cancer Metastasis Rev 1992;11:179–95.

30. Somosy Z., G. Horvath, A. Telbisz, et al. Morphological aspects of ionizing radiation response of small intestine. Micron 2002;33:167–78.

31. Withers H. R., M. M. Elkind. Microcolony survival assay for cells of mouse intestinal mucosa exposed to radiation. Int J Radiat Biol Relat Stud Phys Chem Med 1970;17:261–7.

32. Liu K Palmiero A, Chopra N, Velasquez B, Li Z, Beddar S, Schüler E. Dual beam-current transformer design for monitoring and reporting of electron ultra-high dose rate (flash) beam parameters. Journal of Applied Clinical Medical Physics 2023.

33. Liu K, Holmes S, Hooten B, Schüler E, Beddar S. Evaluation of ion chamber response for applications in electron flash radiotherapy. Medical Physics 2023.

34. Liu K Velasquez B, Schueler E. Technical note: High-dose and ultra-high dose rate (uhdr) evaluation of al2o3:C optically stimulated luminescent dosimeter nanodots and powdered lif:Mg,ti thermoluminescent dosimeters for radiation therapy applications Medical Physics 2023.

35. Liu Kevin, Shannon Holmes, Ahtesham Ullah Khan, et al. Development of novel ionization chambers for reference dosimetry in electron flash radiotherapy. arXiv preprint arXiv:240305661 2024.

36. Baikalov Alexander, Daline Tho, Kevin Liu, et al. Characterization of a novel time-resolved, real-time scintillation dosimetry system for ultra-high dose rate radiation therapy applications. ArXiv 2024.

37. Liu Kevin, Shannon Holmes, Emil Schüler, et al. A comprehensive investigation of the performance of a commercial scintillator system for applications in electron flash radiotherapy. Medical physics 2024.

38. Ashraf M Ramish, Stavros Melemenidis, Kevin Liu, et al. Multi-institutional audit of flash and conventional dosimetry with a 3d printed anatomically realistic mouse phantom. International Journal of Radiation Oncology* Biology* Physics 2024.

39. Palmiero Allison, Kevin Liu, Julie Colnot, et al. On the acceptance, commissioning, and quality assurance of electron flash units. arXiv preprint arXiv:240515146 2024.

40. Valdes Zayas A., N. Kumari, K. Liu, et al. Independent reproduction of the flash effect on the gastrointestinal tract: A multi-institutional comparative study. Cancers (Basel) 2023;15.

41. Rothwell B., M. Lowe, E. Traneus, et al. Treatment planning considerations for the development of flash proton therapy. Radiother Oncol 2022;175:222–230.

42. Diffenderfer E. S., Verginadis, II, M. M. Kim, et al. Design, implementation, and in vivo validation of a novel proton flash radiation therapy system. Int J Radiat Oncol Biol Phys 2020;106:440–448.

43. Zhang Q., E. Cascio, C. Li, et al. Flash investigations using protons: Design of delivery system, preclinical setup and confirmation of flash effect with protons in animal systems. Radiat Res 2020;194:656–664.

44. Levy K., S. Natarajan, J. Wang, et al. Abdominal flash irradiation reduces radiation-induced gastrointestinal toxicity for the treatment of ovarian cancer in mice. Scientific reports 2020;10:21600.

45. Loo B. W., Jr., Verginadis, II, B. S. Sørensen, et al. Navigating the critical translational questions for implementing flash in the clinic. Seminars in radiation oncology 2024;34:351–364.

46. Almeida A., M. Togno, P. Ballesteros-Zebadua, et al. Dosimetric and biologic intercomparison between electron and proton flash beams. Radiother Oncol 2024;190:109953.

47. Gueulette J., V. Gregoire, M. Octave-Prignot, et al. Measurements of radiobiological effectiveness in the 85 mev proton beam produced at the cyclotron cyclone of louvain-la-neuve, belgium. Radiat Res 1996;145:70–4.

48. Tepper J., L. Verhey, M. Goitein, et al. In vivo determinations of rbe in a high energy modulated proton beam using normal tissue reactions and fractionated dose schedules. Int J Radiat Oncol Biol Phys 1977;2:1115–22.

49. Ando K., Y. Furusawa, M. Suzuki, et al. Relative biological effectiveness of the 235 mev proton beams at the national cancer center hospital east. J Radiat Res 2001;42:79–89.

50. Paganetti H., A. Niemierko, M. Ancukiewicz, et al. Relative biological effectiveness (rbe) values for proton beam therapy. Int J Radiat Oncol Biol Phys 2002;53:407–21.

51. Colbert L. E., M. B. El Alam, R. Wang, et al. Tumor-resident lactobacillus iners confer chemoradiation resistance through lactate-induced metabolic rewiring. Cancer Cell 2023;41:1945–1962 e11.

52. Sorensen B. S., E. Kanouta, C. Ankjaergaard, et al. Proton flash: Impact of dose rate and split dose on acute skin toxicity in a murine model. Int J Radiat Oncol Biol Phys 2024.

53. Mascia A., S. McCauley, J. Speth, et al. Impact of multiple beams on the flash effect in soft tissue and skin in mice. Int J Radiat Oncol Biol Phys 2024;118:253–261.

54. Karsch L., J. Pawelke, M. Brand, et al. Beam pulse structure and dose rate as determinants for the flash effect observed in zebrafish embryo. Radiother Oncol 2022;173:49–54.

